# A population phylogenetic view of mitochondrial heteroplasmy

**DOI:** 10.1101/204479

**Authors:** Peter R. Wilton, Arslan Zaidi, Kateryna Makova, Rasmus Nielsen

## Abstract

The mitochondrion has recently emerged as an active player in a myriad of cellular processes. Additionally, it was recently shown that more than 200 diseases are known to be linked to variants in mitochondrial DNA or in nuclear genes interacting with mitochondria. This has reinvigorated interest in its biology and population genetics. Mitochondrial heteroplasmy, or genotypic variation of mitochondria within an individual, is now understood to be common in humans and important in human health. However, it is still not possible to make quantitative predictions about the inheritance of heteroplasmy and its proliferation within the body, partly due to the lack of an appropriate model. Here, we present a population-genetic framework for modeling mitochondrial heteroplasmy as a process that occurs on an ontogenetic phylogeny, with genetic drift and mutation changing heteroplasmy frequencies during the various developmental processes represented in the phylogeny. Using this framework, we develop a Bayesian inference method for inferring rates of mitochondrial genetic drift and mutation at different stages of human life. Applying the method to previously published heteroplasmy frequency data, we demonstrate a severe effective germline bottleneck comprised of the cumulative genetic drift occurring between the divergence of germline and somatic cells in the mother and the separation of germ layers in the offspring. Additionally, we find that the two somatic tissues we analyze here undergo tissue-specific bottlenecks during embryogenesis, less severe than the effective germline bottleneck, and that these somatic tissues experience little additional genetic drift during adulthood. We conclude with a discussion of possible extensions of the ontogenetic phylogeny framework and its possible applications to other ontogenetic processes in addition to mitochondrial heteroplasmy.

## 1. Introduction

As the energy providers of the cell, mitochondria play a vital role in the biology of eukaryotes. Much of the metabolic functionality of the mitochondrion is encoded in the mitochondrial genome, which in humans is ~ 16.5 kb in length and inherited from the mother. While it was long thought that the mitochondria within the human body are genetic clones, it is now recognized that variation of mitochondrial DNA (mtDNA) is common within human cells and tissues. This variation, termed mitochondrial heteroplasmy, is a normal part of healthy human biology (Rebolledo-Jaramillo *et al.*, 2014; Li *et al.*, 2016, 2010), but it is also important in human health and disease, being the primary mode of inheritance of mitochondrial disease and playing a role in cancer and aging (reviewed in Stewart and Chinnery, 2015; Wallace and Chalkia, 2013).

Because of its importance in human health, it is crucial to understand how mitochondrial heteroplasmy is transmitted between generations and becomes distributed within an individual. Heteroplasmy frequencies can change drastically between mother and offspring, owing to a hypothesized bottleneck in the number of segregating units of mitochondrial genomes during early oogenesis (Cree *et al.*, 2008). There has been considerable debate about whether the mechanism of this bottleneck involves an actual decrease in the number of mitochondrial genome copies versus co-segregation of genetically homogeneous groups of mitochondrial DNA (e.g., Jenuth *et al.*, 1996; Cao *et al.*, 2007; Cree *et al.*, 2008; Wai *et al.*, 2008; Carling *et al.*, 2011). Nevertheless, in order to better predict the change in heteroplasmy frequencies between generations, previous studies have sought to infer the size of the oogenic bottleneck, either through direct observation (in mice) of the number of mitochondrial DNA genome copies (Cree *et al.*, 2008; Cao *et al.*, 2007), or through indirect measurement, making statistical conclusions about the bottleneck size based on observed frequency changes between generations (Johnston *et al.*, 2015; Rebolledo-Jaramillo *et al.*, 2014; Millar *et al.*, 2008; Hendy *et al.*, 2009; Li *et al.*, 2016). Recently, Johnston *et al.* (2015) have proposed a statistical framework that combines direct observations of mtDNA copy number with genetic variance in order to make inferences about the dynamics of the oogenic bottleneck. In mice, estimates of the physical bottleneck size have ranged from 200 to more than 1000 (Cree *et al.*, 2008; Cao *et al.*, 2007; Johnston *et al.*, 2015), and in a recent re-analysis of previous data, it was claimed that the minimal bottleneck size may have only small effects on heteroplasmy transmission dynamics, depending on the details of how oogonia proliferate (Johnston *et al.*, 2015). In humans, indirect estimates of the effective genetic bottleneck size have ranged from 1 to 200, depending on the dataset and the statistical methods used to estimate the bottleneck size (Marchington *et al.*, 1997; Guo *et al.*, 2013).

Surveys of heteroplasmy occurrence in humans have also found that heteroplasmic variants are often more numerous and at greater frequency in older individuals, and that older mothers transmit more het-eroplasmies to their offspring (Sondheimer *et al.*, 2011; Rebolledo-Jaramillo *et al.*, 2014; Li *et al.*, 2015). It has also been observed that heteroplasmy frequencies vary from one tissue to another within an individual (Rebolledo-Jaramillo *et al.*, 2014; Li *et al.*, 2015). These observations underscore the fact that heteroplasmy frequencies change not only during oogenesis in the mother, but also during embryogenesis and throughout adult life. Ideally any indirect statistical inferences made about the bottleneck size or other aspects of heteroplasmy frequency dynamics would account for all sources of heteroplasmy frequency change simultaneously. Such an approach would need to account for the phylogenetic and developmental relationships between sampled tissues in order to make full use of the information contained in the observed heteroplasmy allele frequencies. While a number of studies have employed or developed population-genetic models to study mitochondrial heteroplasmy (e.g., Wonnapinij *et al.*, 2008; Hendy *et al.*, 2009; Johnston *et al.*, 2015; Johnston and Jones, 2016), few have considered the phylogenetic relationship between tissues in doing so, often because only a single tissue type is under consideration. Recently, Burgstaller *et al.* (2014) inferred tissue-specific rates of heteroplasmy segregation in artificially heteroplasmic mouse lines, implicitly relating the sampled tissues by a star-like phylogeny. In a study of heteroplasmy frequencies sampled from 11 tissues in unrelated individuals, Li *et al.* (2015) constructed a phylogeny of the tissues but did not combine it with population-genetic analysis.

Here, we describe a model of heteroplasmy dynamics throughout several key stages of human growth and reproduction. Our approach is to model heteroplasmy frequency change as a population-genetic process of genetic drift and mutation that occurs along the branches of an ontogenetic phylogeny, which we define as the tree-like structure relating sampled tissues by their developmental and ontogenetic histories. Our model is similar to typical population-phylogenetic inference models (e.g., Pickrell and Pritchard, 2012; Gautier and Vitalis, 2013), but it also includes features that are unique to ontogenetic phylogenies. We employ our model in a Bayesian inference procedure that uses Markov chain Monte Carlo (MCMC) to sample from posterior distributions of genetic drift and mutation rate parameters for various developmental processes. After demonstrating the accuracy of our method with simulated data, we apply it to real heteroplasmy frequency data and present new insights into the dynamics of heteroplasmy frequency change in humans.

## 2. Methods

### 2.1. Ontogenetic phylogenies

We model the mitochondria in tissues sampled from one or more related individuals as a group of populations related by an ontogenetic phylogeny. Along each branch of the ontogenetic phylogeny, heteroplasmy frequencies within some ancestral tissue change due to the action of genetic drift and mutation. We assume that the shape of the ontogenetic phylogeny is given.

Our ontogenetic phylogeny model differs in a few important ways from the typical population-phylogenetic likelihood framework. In the typical population-genetic model, each branch is considered to be an independent period of evolutionary history and thus is under control of an independent parameter. In contrast to this, we allow a single parameter to determine the dynamics on multiple parts of the phylogeny, since a single developmental process can act in multiple related individuals, and this developmental process can be assumed to act similarly in each individual. Furthermore, while it is typically assumed that each locus has been transmitted through a single phylogeny and thus has been subject to the same population-genetic forces, we allow the effects of genetic drift and mutation to depend on the age of the sampled individuals. In particular, for certain ontogenetic processes, we model the rate of accumulation of genetic drift and mutation with age. This is motivated by previous observations that heteroplasmic variants segregate and accumulate with time within somatic tissues (Li *et al*, 2015; SONDHEiMER *et al.*, 2011; REBOLLEDO-JARAMiLLO *et al*, 2014) and within the germline (REBOLLEDO-JARAMiLLO *et al.*, 2014; Li *et al.*, 2016; WACHSMUTH *et al.*, 2016). Finally, in the typical population-phylogenetic model, each branch of the phylogeny represents a single period in evolutionary history and is modeled by a single parameter. Because multiple ontogenetic processes of interest can occur along a single branch of an ontogenetic phylogeny, we allow branches on the ontogenetic phylogeny to be broken into multiple distinct ontogenetic processes, controlled by independent parameters. Figure 1 demonstrates these features with an ontogenetic phylogeny representing the relationships between two tissues sampled in both a mother and her offspring.

**Figure 1:**
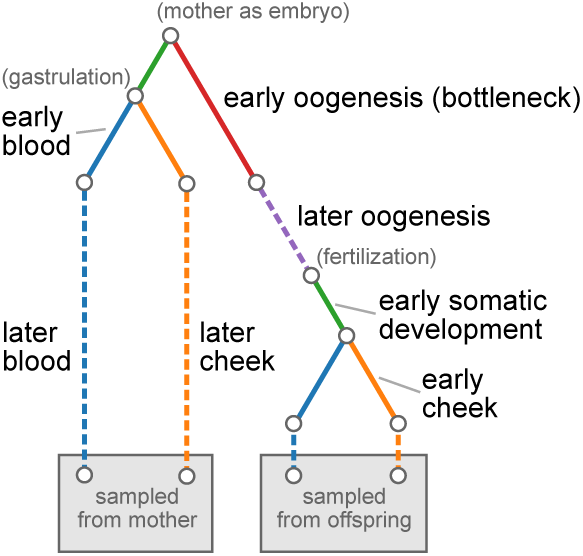
Ontogenetic phylogeny for sampled tissues in mother-child duos from REBOLLEDO-JARAMILLO *et al.* (2014). Each color represents a different tissue or developmental process. The leaves of the tree represent the blood and cheek epithelial tissues sampled from the mother and her child. Solid lines show processes modeled by a fixed amount of genetic drift and dashed lines show processes in which genetic drift accumulates linearly with age. The red component, representing early oogenesis, models a single-generation bottleneck with subsequent doubling of the population size back up to a large size. Parenthetical descriptions in gray show the timing of notable developmental events.

Each ontogenetic process in the phylogeny is parameterized by a genetic drift parameter and a mutation rate. The mutation rate is *θ* = 2*N*_*eμ*_, where *N*_*e*_ is the effective size of the relevant cell population and μ is the per-replication, per-base mutation rate. Genetic drift can be modeled in one of three ways, namely as a fixed amount of genetic drift, as an explicit bottleneck size, or as a rate of accumulation of genetic drift per year.

### 2.2. Likelihood calculation

Given ontogenetic tree *T* with *k* ontogenetic processes, genetic drift parameters *b* = {*b*_1_,…, *b*_*k*_} and mutation rates *θ* = {*θ*_1_,…, *θ*_*k*_}, our likelihood is

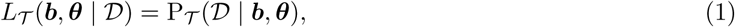

where *D* represents the heteroplasmy frequency data. (Below, the *T* subscript is left off for brevity.) Suppose heteroplasmy frequencies were sampled from *F* families. Writing *C*_*i*_ for the number of heteroplasmic sites in family *i*, *D*_*ij*_ for the heteroplasmy frequency data at the *j*th heteroplasmic locus (of *C*_*i*_) in family *i*, and *H*_*ij*_ for the event that site *j* is heteroplasmic in family *i*, our likelihood can be written

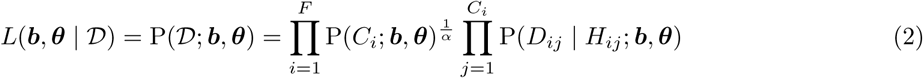

where P(*C*_*i*_; *b, θ*) is the probability of *C*_*i*_ heteroplasmic variants occurring in family *i* and P(*D*_*ij*_ | *H*_*ij*_; *b, θ*) is the probability of the observed heteroplasmy data at the *jth* heteroplasmic locus in family *i*, conditional on heteroplasmy (i.e., polymorphism) in at least one tissue at that locus. We assume that C_i_ is Poisson distributed with rate *G* · *P*(*H*_*ij*_; *b, θ*), where *G* is the genome size and P(*H*_*ij*_; *b, θ*) is the probability that site *j* is heteroplasmic in family *i*. We note that P(*H*_*ij*_) = P(*H*_*ik*_) for each *j* and *k*; that is, the probability of heteroplasmy depends on the family (specifically, on the age of individuals in the family) and not on the particular locus.

We penalize the part of the likelihood involving the number of heteroplasmic variants with the parameter *α* in order to make inference less sensitive to experimental heteroplasmy detection, which is a non-trivial problem, especially for heteroplasmies segregating at low frequency (Li and STONEKiNG, 2012; REBOLLEDO-JARAMiLLO *et al*, 2014). Without such a penalty, the likelihood is too strongly influenced by the number of observed heteroplasmies, a quantity influenced both by false positives—at a rate of up to ~10% for low-frequency heteroplasmies in REBOLLEDO-JARAMiLLO *et al.* (2014)—and by false negatives caused by conservative minimum allele frequencies thresholds (1% in REBOLLEDO-JARAMiLLO *et al.*, 2014). On the other hand, if the number of heteroplasmies is completely absent from the likelihood, such that all information about drift and mutation is taken only from the heteroplasmy frequencies, posterior distributions of mutation rates are sensitive to outlier allele frequencies that do not fit a model of genetic drift and (infrequent) mutation as well. As a compromise, we set the value of this likelihood penalty to *α* = 100, which in effect artificially reduces the total number of sites considered in this component of the likelihood, such that if in reality 500 heteroplasmic sites are observed out of a total of 100, 000 sites, the contribution to the likelihood would be the same as if 5 heteroplasmic sites were observed in a total of 1000 sites.

With our likelihood (2) we implicitly ignore linkage between heteroplasmic sites within a family even though in reality the lack of recombination means that the sites are perfectly linked. We justify this approximation in two ways: first, there are usually few heteroplasmic variants co-segregating in a family (mean 2.6 in REBOLLEDO-JARAMiLLO *et al.* 2014, 1.0 in Li *et al.* 2016), and second, amongst heteroplasmic variants co-segregating in a family, most segregate at low frequency, so that changes in the frequency of one heteroplasmy do not greatly affect the frequency of another. Thus the dynamics at several heteroplasmic sites should closely resemble those of a model in which each site truly segregates independently. This assumption is supported by simulations of nonrecombining mitochondrial genomes (see Section 2.4 below). We further assume that heteroplasmy frequencies are independent between families.

A site is determined to be heteroplasmic according to the filtering steps described in REBOLLEDO-JARAMiLLO *et al.* (2014), which include filters for mapping quality, base quality, minimum allele frequency (1%), coverage (> 1000x), local sequence complexity, and contamination. Rather than calculate likelihoods based on called allele frequencies, we model binomial sampling error in the number of consensus and alternative reads sampled from a true, unknown allele frequency. Thus *D*_*ij*_ represents the number of consensus and alternative alleles at the jth heteroplasmic locus in family *i*. Conditional on heteroplasmy (i.e., polymorphism), the probability of the observed read counts *D*_*ij*_ at locus *j* in family *i* is

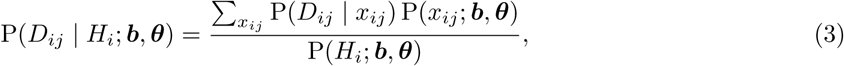

where *x*_*ij*_ is the true, unknown allele frequency at locus *j* in family *i*. The sum is performed over all possible allele frequencies in the sampled tissues. Both the numerator and the denominator can be calculated using FELSENSTEiN’s (1981) pruning algorithm, a dynamic programming algorithm frequently used in likelihood calculations for phylogenetic trees. Details of how we calculated these quantities are given in Appendix A. The pruning algorithm requires calculating allele frequency transition distributions for different genetic drift and mutation parameter values. As described in Appendix B, we achieved this by numerically pre-computing allele frequency transition distributions under the discrete-generation Wright-Fisher model and linearly interpolating between pre-computed values. The pruning algorithm also requires a distribution of allele frequencies at the root of the phylogeny, which, in our application (see below), represents the unobservable distribution of heteroplasmy allele frequencies in the mother as an embryo. Following TATARU *et al.* (2015), we use a discretized, symmetric beta distribution with additional, symmetric probability weights at frequencies 0 and 1. The two parameters specifying this distribution are inferred jointly with genetic drift and mutation parameters.

### 2.3. Inference

We take a Bayesian approach to inference. Prior distributions are Log-Uniform(5 × 10^−4^,3) for genetic drift parameters, measured in generations per *N*_*e*_ (henceforth “drift units”). For genetic drift parameters specified by a rate of accumulation of drift units per year, the lower (resp. upper) limit of the (Uniform) prior distribution limits are divided by the minimum (resp. maximum) of the ages by which the rate is multiplied. We did not allow the effects of genetic drift to decrease with age. Prior distributions on bottleneck sizes are Log-Uniform(2, 500), and for mutation rate parameters *θ* = 2*N*_*eμ*_, the prior distribution is Log-Uniform(10^−8^,10^−1^).

We employ an affine-invariant ensemble Markov Chain Monte Carlo (MCMC) procedure (Goodman and Weare, 2010) to sample from posterior distributions, as implemented in the Python package emcee (Foreman-Mackey *et al.*, 2013). We assess convergence by visual inspection of the posterior traces. Running 500 chains in the ensemble MCMC for 20000 iterations each, we find good convergence after ~2500 iterations and thus discard the first 5000 iterations of each chain as burn-in. With ~100 heteroplasmic loci, a run takes 60–80 CPU hours, but due to the parallel nature of ensemble MCMC, calculations can be efficiently spread across CPUs, so that on a twenty-core compute node, results are obtained in approximately four hours. Reported 95% credible intervals are intervals of the highest posterior density.

As a way of evaluating the relative support for different ontogenetic models, we estimate Bayes factors (i.e., ratios of posterior evidence integrals) for alternative ontogenetic models of the accumulation of drift within cell lineages. For models *M*_1_ and *M*_2_, the Bayes factor is

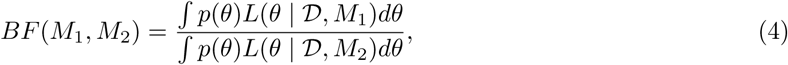

wherep(·) is the prior distribution and *L*(· | *D*, *M*_*k*_) is the likelihood under model *k*. These posterior evidence integrals are approximated using emcee’s (Foreman-Mackey *et al.*, 2013) implementation of an approach using thermodynamic integration (see Goggans and Chi, 2004).

### 2.4. Simulation

We performed two sets of simulations to test our inference procedure. The first simulations were performed under the model assumed by our inference procedure. As described above, this model assumes that each locus segregates independently, allele frequency transitions occur according to the Wright-Fisher model of genetic drift and bi-allelic mutation, and heteroplasmy frequencies in the root of the ontogenetic phylogeny are controlled by the two parameters of a discretized, symmetric beta distribution with extra probability weight at frequencies zero and one. These simulations were performed forward in time using a custom Python script.

The second set of simulations tested how our assumption that loci segregate independently affects inference when the data are simulated from nonrecombining genomes sampled from many different families. These simulations were performed using a custom interface to the simulation package msprime (Kelleher *et al.*, 2016), which simulates genetic variation under the standard neutral coalescent model with infinite-sites mutation. In these simulations, population sizes and branch lengths are equivalent to those under the forward-time simulations, but at the root of the ontogenetic phylogeny, we assume that ancestral lineages trace their ancestry back in time in a single panmictic population of constant size. Simulations were performed under conditions in which the distribution of the number of heteroplasmic variants per family roughly matched the distribution observed in the data.

### 2.5. Data

We applied our inference procedure to a publicly available dataset, containing allele frequencies for 98 het-eroplasmies sampled from from 39 mother-offspring duos, originally published by Rebolledo-Jaramillo *et al.* (2014). In this dataset, mitochondria from blood and cheek epithelial cells were sampled from both mother and offspring, resulting in a ontogenetic phylogeny with four leaves, each representing one of the four tissues sampled from a mother-offspring duo. Details of heteroplasmy discovery are described in Rebolledo-Jaramillo *et al.* (2014).

To model the segregation of heteroplasmy frequencies during the ontogeny of the four tissues sampled from each duo, we used the ontogenetic phylogeny shown in Figure 1. This ontogenetic phylogeny models several life stages. The root of the phylogeny occurs at the divergence of the mother’s somatic and germline tissues when she is an embryo. On the branch leading to the somatic tissues in the mother, there is a brief period of early embryonic development before the blood and cheek epithelial cell lineages diverge at gastrulation as members of the ectodermal (cheek epithelial) and mesodermal (blood) germ layers. After diverging at gastrulation, each somatic tissue undergoes independent periods of genetic drift and mutation during later embryogenesis and early growth, and finally for each tissue there are independent rates of accumulation of genetic drift and mutation throughout adult life.

On the branch leading to the offspring tissues in the ontogenetic phylogeny in Figure 1 the first stage represented is the period of oogenesis prior to the birth of the mother, when the oogenic bottleneck is thought to occur. This is followed by the oocyte stage, during which we assume the mitochondria accumulate genetic drift and mutation at some rate linearly with the age of the mother before childbirth. At fertilization, this branch undergoes the same period of early somatic development experienced by the mother’s somatic tissues prior to gastrulation. Finally, the two somatic tissues of the offspring diverge at gastrulation and go through the same stages of development as the somatic tissues of the mother. For an overview of the events of human development, see, for example, Carlson (2014).

### 2.6. Effective oogenic bottleneck

Analyzing both simulated and real data, we find that there is limited power to infer the size of the oogenic bottleneck. This is to be expected, given that we also model the subsequent genetic drift of the later stages of oocyte development and in the early developing embryo; each of these three ontogenetic processes occurs along the same branch of the ontogenetic phylogeny of the tissues considered here (Fig. 1), which causes their respective contributions of genetic drift to be conflated with one another. We note that the genetic drift parameters of these ontogenetic processes are not truly unidentifiable: power to distinguish genetic drift during the early-oogenesis bottleneck from that of the later maternal germline is provided by the differing effects of genetic drift in mothers of different ages, and power to distinguish the contribution of drift in the early embryo is provided by the fact that this process occurs in both the mother and the offspring. Differences in effective population size (and thus scaled mutation rates) also provide theoretical power to distinguish these parameters, but nevertheless we find that these genetic drift parameters tend to become conflated with one another.

As a way of counteracting this conflation, we combine the genetic drift parameters of this branch in the ontogenetic phylogeny into an effective bottleneck size (EBS), summarizing the total genetic drift between mother and offspring. The effective bottleneck is comprised of the oogenic bottleneck per se, the accumulation of genetic drift in the oocyte prior to ovulation, and the genetic drift in the embryo between fertilization and gastrulation. To combine genetic drift parameterized as a bottleneck with genetic drift parameterized in drift units, we used the approximate relationship *N*_*b*_ ≈ 2/*d*, where *d* is genetic drift in drift units, and *N*_*b*_ is the bottleneck size. This approximation is justified in Appendix C. Using this relationship, our equation for the EBS has the form

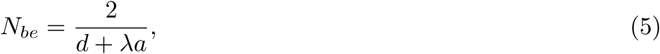

where *d* is the summed genetic drift from the oogenic bottleneck *per se* and pre-gastrulation embryogenesis, *λ* is the rate of genetic drift accumulation in the oocyte, and *a* is the age of the mother at childbirth. Because in our model genetic drift accumulates in the oocyte as the mother ages prior to ovulation, the size of the effective bottleneck decreases with age. We summarize this rate of decrease by linearizing (5) between ages 25 and 34, the first and third quartiles of maternal age at childbirth in the dataset from Rebolledo-Jaramillo *et al.* (2014).

### 2.7. Availability

Our inference procedure is released under a permissive license in a Python package called mope, available at https://github.com/ammodramus/mope or from the Python Package Index (PyPI, http://pypi.python.org/). As we describe above, our inference procedure requires precomputed transition distributions. These can be generated by the user or downloaded from https://github.com/ammodramus/mope. Our simulation scripts are also provided with the inference procedure.

Data from Rebolledo-Jaramillo *et al.* (2014) are available from that paper’s supplementary material and from the NCBI Sequence Read Archive (www.ncbi.nlm.nih.gov/sra), accession SRP047378.

## 3. Results

### 3.1. Application to simulated data

The targets of our inference procedure are genetic drift parameters and population-size-scaled mutation rates for each ontogenetic process in the ontogenetic phylogeny. Genetic drift may be parameterized as a fixed amount of genetic drift (in drift units, i.e. generations / *N*_*e*_), as a rate of accumulation of drift per year, or as a haploid bottleneck size. The scaled mutation rates, *θ* = 2*N*_*eμ*_ are twice the product of the haploid effective population size *N*_*e*_ and the per-replication, per-base mutation rate *μ*. Since *μ* can be assumed to be the same in every mitochondrion, the mutation rates can also be interpreted as relative effective population sizes. Two parameters controlling the distribution of allele frequencies at the root of the phylogeny are also inferred.

The inference procedure performed well on data simulated under the model of drift and mutation assumed by the inference procedure. In a simulation of 500 independently segregating sites sampled from two tissues in each of 100 different mothers and their offspring, under parameters producing a total of 110 heteroplasmic variants, the branch lengths and mutation rates were inferred without apparent bias (Fig. 2), as were the two root distribution parameters (not shown). Posterior distributions were generally narrower for genetic drift parameters than for scaled mutation rates, likely corresponding to the fact that there is less information about mutation than genetic drift in the simulated data. Parameters of external branches were inferred more precisely than those of internal branches. Other parameter values produced similar results (Fig. S1).

**Figure 2:**
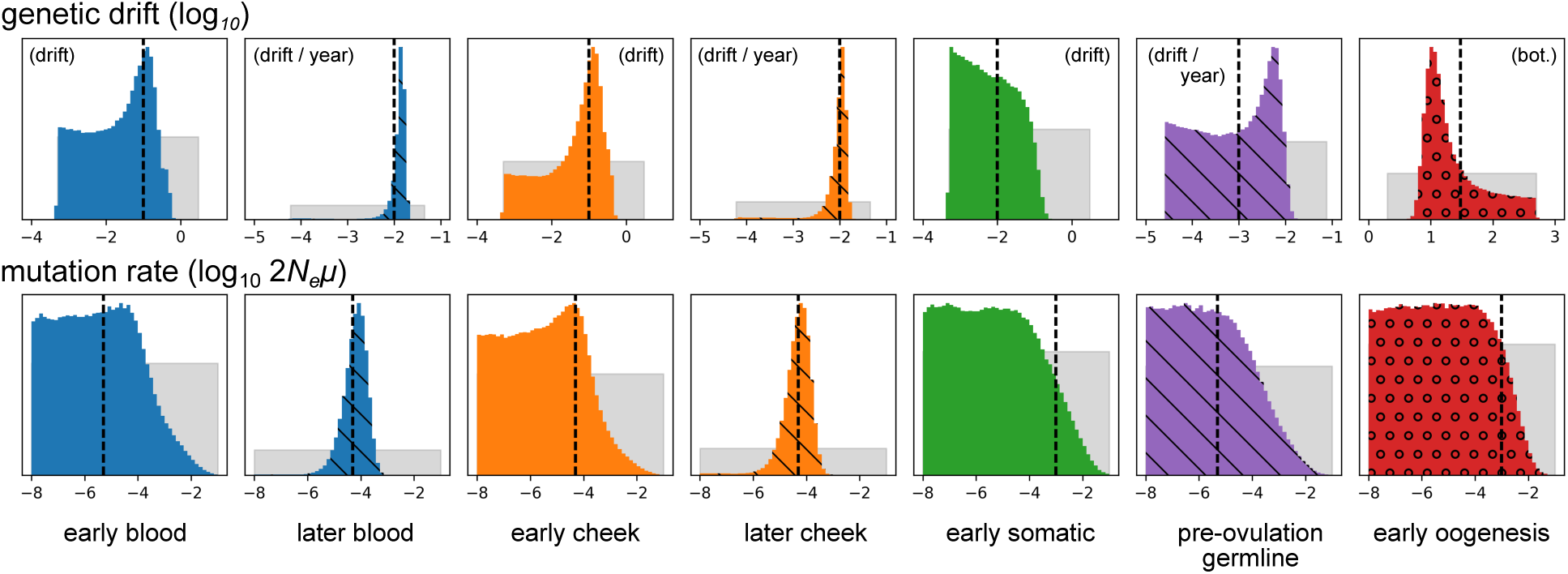
Posterior distributions of genetic drift and mutation parameters inferred from data simulated under the model assumed by the inference procedure. The top and bottom rows depict genetic drift and mutation rate parameters, respectively. Gray distributions depict prior distributions, and colored distributions depict posterior distributions. Colors match the colors of the ontogenetic processes in Figure 1. Distributions hashed with diagonal lines correspond to processes with drift parameterized by rates of accumulation of genetic drift with age. (That is, they correspond to the dashed lines in Fig. 1.) The circles in the red posterior distributions indicate that this process is modeled by an explicit bottleneck. All parameters are log_10_-transformed, and the distributions correspond to these transformed variables. Vertical dashed lines show the simulated parameter values. Not shown are the two parameters controlling the allele frequency distribution at the root of the phylogeny, which were inferred with comparable accuracy.

The procedure also performed well on data generated in simulations that did not assume free recombination between heteroplasmic sites (Fig. S2). In these simulations, we simulated non-recombining mitochondrial genomes of 10, 000 base pairs in 30 mother-offspring duos, under parameters resulting in 104 heteroplasmic variants. The ~3.7 heteroplasmies per family in these simulations is similar to the ~2.6 observed in the data from Rebolledo-Jaramillo *et al.* (2014), supporting our assumption that linkage between heteroplasmic variants within families does not greatly affect inference results. We also tested for robustness against false positive and false negative heteroplasmy detection. Applying our method to simulations with a false negative rate of 0.4 for truly heteroplasmic mutations between frequencies 0.1% and 2%, and a false positive rate of 3 × 10^−5^ per bp, so that 14 false negatives and 5 false positives were produced in a dataset of 101 heteroplasmic loci, the inference procedure was not apparently biased away from the true, simulated parameter values (Fig. S3).

### 3.2. Application to real heteroplasmy data

In the application of our method to the heteroplasmy frequency data from Rebolledo-Jaramillo *et al.* (2014) (Fig. 3), we find that the posterior distribution of the size of the early oogenesis bottleneck is broad, with a 95% credible interval (CI) spanning from 10.6 to 433.2. As we describe above (see 2.6), this is unsurprising given that in the assumed ontogenetic phylogeny there are three independent periods of drift and mutation along the branch containing the oogenic bottleneck, namely the early oogenic bottleneck itself, the turnover of mitochondria in the oocyte prior to ovulation, and the period after fertilization but before gastrulation (Fig. 1).

**Figure 3:**
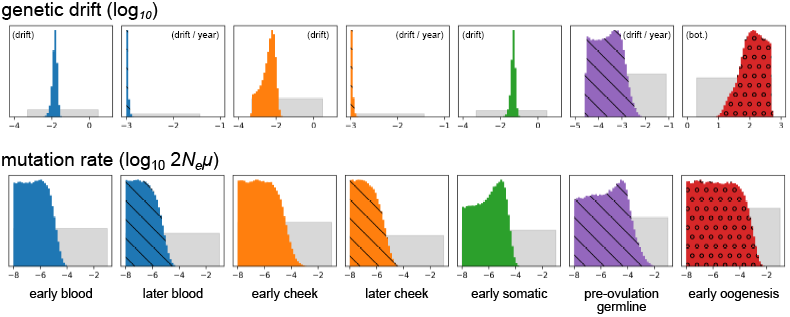
Inference results for real heteroplasmy frequency data. The top row shows results for genetic drift parameters, and the bottom row shows posterior distributions for scaled mutation rates. Distributions hashed with diagonal lines correspond to processes with drift parameterized by rates of accumulation of genetic drift with age. (That is, they correspond to the dashed lines in Fig. 1.) The circles in the red posterior distributions indicate that this process is modeled by an explicit bottleneck. All parameters are log_10_-transformed, and the depicted distributions correspond to these transformed variables. Distributions are not drawn to a common vertical axis.

To counteract this conflation, we combined the genetic drift into an effective bottleneck. The posterior distribution of the size of this effective bottleneck (i.e., the EBS) was substantially narrower than that of the explicitly modeled bottleneck, with a median of 24.5 (11.6–35.1, 95% CI) for a mother of the mean age in this dataset (Fig. 4A). This is in line with the bottleneck size estimate of 32.3 produced by Rebolledo-Jaramillo *et al.* (2014).

**Figure 4:**
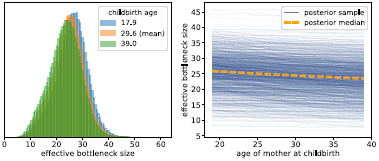
Posterior samples of the effective bottleneck size for mothers of different ages. (A) Posterior distribution of the effective between-generation bottleneck size for younger, older, and median-aged mothers. (B) Relationship between mother’s age at childbirth and the effective oogenic bottleneck size. The orange dashed line shows how the median effective bottleneck size varies with age at childbirth. The solid blue lines show posterior samples from the relationship between effective bottleneck size and age at childbirth, with each having the form of (C.4), where the genetic drift parameters in this equation are jointly sampled from the posterior distribution. A total of n = 1000 lines sampled from the posterior are plotted. We note that each line necessarily decreases with mother birth age due to our assumption that genetic drift accumulates at some rate in the oocyte (see (C.4)); what varies from one line to another is the rate at which the effective bottleneck size decreases due to this accumulation of genetic drift.

In our model, genetic drift accumulates in the oocyte as the mother ages, and thus the size of the effective bottleneck decreases with age of the mother at childbirth. The inferred relationship between age at childbirth and EBS is shown in Figure 4B. At age 18, the median posterior EBS is 26.1 (13.0–36.9, 95% CI), and at age 40, it is 23.4 (10.5–34.2). The median posterior rate of decrease of the EBS is −0.075 bottleneck units per year, although the central 95% credible interval for this rate of decrease is broad (0.0–0.34). Given the range of this credible interval, there is apparently limited information contained in the data about whether or not the EBS decreases with age, or equivalently, whether genetic drift accumulates meaningfully in the oocyte.

The median posterior rates of genetic drift accumulation in adult somatic tissues were very small, just 1.0 × 10^−3^ (1.0 × 10^−3^–1.2 × 10^−3^, 95% CI) drift units per year for blood, and 1.0 × 10^−3^ (1.0 × 10^−3^– 1.2×10^−3^) drift units per year for cheek. These estimates are at the lower limit of what is permissible under our model of genetic drift, which is based upon distributions of allele frequency change in a finite-sized Wright-Fisher model (see Appendix B). On the other hand, the inferred amounts of genetic drift occurring during early development of the somatic tissues was greater: 0.015 (0.0067–0.023, 95% CI) drift units for blood, and 0.0044 (5.0 × 10^−4^–0.011) drift units for cheek, roughly equivalent to bottlenecks of size 136.2 (74.1–247.5) and 457.7 (103.9–2817.1), respectively.

The posterior distributions of scaled mutation rates were broad, and thus limited information about the relative population sizes of different developmental and adult tissues is contained in the heteroplasmy frequency data. This is unsurprising given that the problem is similar to attempting to infer population size history from ~100 single-nucleotide polymorphisms. A high scaled mutation rate (2*N*_*eμ*_ > 10^−4^) is (relatively) most supported in oogenesis, reflecting the observation of possibly *de novo* mutations in the dataset. However, the 95% credible interval of each developmental process spans several orders of magnitude (at least 10^−8^ < 2*N*_*eμ*_ < 10^−5^), so firm conclusions cannot be drawn.

We assessed the fit of our model to the real heteroplasmy data by simulating data under the maximum *a posteriori* (MAP) parameter values and comparing to the real data. Comparing the marginal distribution of allele frequencies in the sampled tissues (i.e., the marginal site-frequency spectrum) from the actual data to the MAP simulation data, we find that the marginal distribution of allele frequencies is similar between the two datasets (Fig. 5A), as is the distribution of absolute differences between each pair of sampled tissues (Fig. 5B).

**Figure 5:**
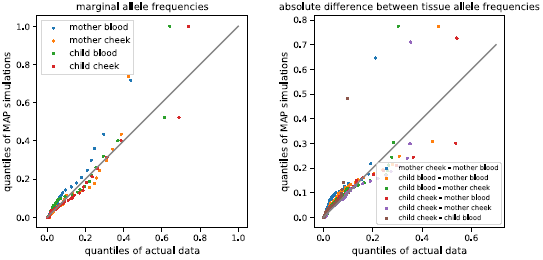
Quantile-quantile comparison of real heteroplasmy data from Rebolledo-Jaramillo *et al.* (2014) and data simulated under maximum *a posteriori* parameter estimates inferred from this data. Panel (A) compares marginal distributions of allele frequencies in each tissue, and panel (B) compares distributions of absolute differences in allele frequency between tissues. Each dot represents a sequential percentile of the distributions being compared. Following Rebolledo-Jaramillo *et al.* (2014), alleles were polarized such that the minor allele in the mother (averaged across her two tissues) was denoted as the focal allele.

In order to use Bayes factors (4) to compare the support for different ontogenetic phylogenies, we calculated the posterior evidence integral for the ontogenetic phylogeny in Figure 1 as well as for two additional ontogenetic phylogenies differing in their assumptions about how genetic drift accumulates in somatic tissues (Fig. S4). The first additional model (termed “fixed”, Fig. S4A), assumes that all genetic drift and mutation particular to each somatic tissue occurs early during development and that there is no additional drift accumulating later in life. The second, (“linear”, Fig. S4B), assumes that genetic drift and mutation accumulate linearly with age in somatic tissues. Our original model (Fig. 1) we term “both”, since it assumes that genetic drift both occurs in a fixed quantity during early development and accumulates later in life.

We find that the “fixed” model is more supported than the “both” or “linear” models, with the approximate log-evidence values of the “fixed”, “both”, and “linear” models being –1704±3, –1764±4, and –1816 ± 3, respectively. In the “both” model, in which there is both a period of genetic drift and mutation in the somatic tissues during early development, the inferred rates of drift accumulation are at the minimum allowed by the inference method (~ 10^−3^ drift units per year). This, together with the fact that the best-supported model does not include the accumulation of genetic drift in adult somatic tissues, suggests that there is very little additional genetic drift occurring after birth in the two somatic tissues considered here.

## 4. Discussion

Because we modeled genetic drift during multiple ontogenetic processes between embryogenesis in the mother and the sampling of tissues in the child, our estimate of the size of the oogenic bottleneck per se was imprecise, with a broad 95% credible interval (10.6–433.1). This is concordant with a recent analysis of the time-evolution of heteroplasmy variance in mouse oocytes, which concluded that the actual minimal bottleneck size is difficult to determine and may have only limited impact on overall heteroplasmy dynamics during oogenesis (Johnston *et al.*, 2015). However, our estimates of the EBS (median 24.5, 95% CI: 11.6– 35.1) are similar to other recent estimates of the oogenic bottleneck size, including an estimate of 32.3 in a previous analysis of the data used in this study (Rebolledo-Jaramillo *et al.*, 2014), and a previous estimate of 9 in Li *et al.* (2016).

Our inference framework allows for the size of the effective oogenic bottleneck to decrease with the age of the mother as genetic drift accumulates in the oocyte. We found a broad posterior distribution of the rate by which the EBS decreases in the oocyte (roughly 0.00–0.34 bottleneck units per year, 95% CI), demonstrating that with the 39 mother-child pairs and 98 heteroplasmic variants in the dataset we analyzed (Rebolledo-Jaramillo *et al.*, 2014), there is insufficient information obtained by our model to determine whether genetic drift accumulates with age in the oocyte. In the future, sampling more individuals and tissues, and with larger pedigrees, it may be possible to provide stronger statistical evidence for or against genetic drift occurring in the oocyte; this will potentially be informative on the question of how mitophagy and mitochondrial turnover are involved in oocyte aging, a topic of interest in the study of human fertility (see Zhang *et al.*, 2017).

In addition to the effective bottleneck between mother and offspring, we also quantified genetic drift occurring during the embryonic development of the blood and cheek epithelial lineages. We found that the embryonic genetic drift of heteroplasmy frequencies specific to these tissues was less than the effective between-generation bottleneck but still appreciable, with median posterior estimates of the effective bottleneck sizes being 136.2 (74.1–247.5, 95% CI) and 457.7 (103.9–2817.1) for blood and cheek epithelial cells, respectively.

At the same time we inferred that there is little accumulation of genetic drift in adult somatic tissues. This may seem to contradict previous observations that heteroplasmies become more numerous with age (e.g., Rebolledo-Jaramillo *et al.*, 2014; Li *et al.*, 2016). If the effective population size of the somatic stem cells supporting mitotic somatic tissues is larger than the effective population size during embryogenesis or the maternal germ line, an accumulation of genetic drift with age would produce additional de novo somatic heteroplasmies. On the other hand, if effective population sizes of somatic stem cells are smaller than effective population sizes during early development, a longer period of genetic drift in adulthood would result in fewer heteroplasmic loci, as genetic variation is lost due to ongoing genetic drift in a smaller population. Here, the posterior distributions of population-scaled mutation rates are too broad to permit anything to be concluded about the relative sizes of relevant stem cell populations.

There are several ways our inference procedure could be extended. Our model assumes selective neutrality, but it is possible, or even likely, that neutral population-genetic models do not completely describe the dynamics of heteroplasmy frequency change. Studies of heteroplasmy occurrence in humans have found a relative lack of non-synonymous heteroplasmic mutations (Ye *et al.*, 2014; Rebolledo-Jaramillo *et al.*, 2014), or an excess of non-synonymous mutations at low versus high frequencies (Li *et al.*, 2016), suggesting purifying selection. However, evidence for biased transmission of the major heteroplasmic allele over the minor allele has been inconsistent, with one recent study finding no systematic difference in heteroplasmy allele frequency between other offspring (Li *et al.*, 2016), while the original publication of the data analyzed here did find transmission to be biased towards the major allele at non-synonymous sites (Rebolledo-Jaramillo *et al.*, 2014). Other studies have also found evidence for positive selection acting on heteroplasmies in somatic tissues, observing repeated occurrence of tissue-specific and allele-specific heteroplasmies in many unrelated individuals (Samuels *et al.*, 2013; Li *et al.*, 2015). Studies in mice have also indicated that heteroplasmy may be under natural selection in many instances (e.g., Fan *et al.*, 2008; Stewart *et al.*, 2008; Sharpley *et al.*, 2012; Burgstaller *et al.*, 2014).

It is possible that the systematic biases in model fit represented in Figure 5 are caused by unaccounted-for natural selection. For example, compared to the observed distribution of heteroplasmy frequencies, the MAP model parameters produce an overabundance of intermediate-to-high-frequency heteroplasmies in blood tissues (Fig. 5). Hypothetically, this could be caused by purifying selection against harmful heteroplasmic mutations in blood, which could skew the distribution of heteroplasmy frequencies towards zero. If selection tends to act on only a single heteroplasmic variant at a given time (i.e., if clonal interference between different heteroplasmic alleles is rare), the method we present here could potentially be adapted to make inferences about natural selection in place of mutation. We leave this for future work.

We note that in a recent study finding repeated convergent heteroplasmy in specific tissues in humans, and thus evidence of positive selection on heteroplasmy (Li *et al.*, 2015), the subjects under consideration were deceased and thus older than those considered by Rebolledo-Jaramillo *et al.* (2014); if selection on mitochondrial heteroplasmy intensifies with age, this may explain the lack of such repeated convergence in Rebolledo-Jaramillo *et al.* (2014).

We chose to model heteroplasmy allele frequency dynamics with the Wright-Fisher population model from population genetics. This model is well-studied and thus facilitates interpretation, and it is general in the sense that many different population-genetic models of reproduction closely resemble the Wright-Fisher model when population sizes are at least moderately large (Ewens, 2004). The Wright-Fisher model does not include mechanistic details of mtDNA dynamics such as the hypothesized segregation of mtDNA copies in genetically homogeneous nucleoids (e.g., Cao *et al.*, 2007; Khrapko, 2008), or mitochondrial fission, fusion, degradation, and duplication. The coarse effects of many of these mechanistic details are likely to be captured by the Wright-Fisher model through appeals to the concept of an effective population size, just as the details of reproduction of many classical models of reproduction from population genetics can often be reduced to a change in the effective population size of the Wright-Fisher model (Ewens, 2004; Wakeley, 2009). Here, if we were to include these effects in our model, there would likely be very little power to infer their properties, as sample sizes are small (n < 100 heteroplasmies). In larger and differently structured datasets, there may be greater power to infer mechanistic details of mitochondrial proliferation.

In a study of mitochondrial heteroplasmy transmission between the mothers and children of two-parent-child trios from the Netherlands, Li *et al.* (2016) found support for a variable bottleneck size, where the size of the bottleneck for a particular heteroplasmic locus is randomly sampled from a distribution. The model we present here also allows for variable bottleneck sizes, but it assumes a particular relationship between the effective oogenic bottleneck size and the age of the mother. As discussed above, our inference is inconclusive about whether or not the bottleneck size is variable with age. A variable bottleneck size, independent of mother’s age, could also be implemented in our inference framework by integrating over the distribution of bottleneck sizes during the calculation of allele frequency transition distributions. In this case, like Li *et al.* (2016), we would be inferring the parameters of the bottleneck size distribution rather than a single bottleneck size. We leave this as an opportunity for future investigation.

Johnston *et al.* (2015) have recently used a detailed, mechanistic model of mitochondrial duplication, degradation, and partitioning to study mitochondrial dynamics during oogenesis. The authors applied their model to data on the time evolution of heteroplasmy frequency variance and mtDNA copy number variation during oogenesis in mice, finding that the size of the oogenic bottleneck is just one contributor to the final variance in heteroplasmy frequencies after oogenesis is complete, and that their analysis is inconclusive about the fine details of segregation in nucleoids (except that nucleoids are not very large and genetically homogeneous). This work is broadly in agreement with the present study and is complementary in that it analyzes just one phase of ontogeny (namely, oogenesis) and makes use of time series observations of heteroplasmy frequencies in mice rather than heteroplasmy frequencies in multiple somatic tissues in adult humans.

However, it is still possible that the dynamics of heteroplasmy frequency change do not meet the basic assumptions of any population-genetic model. Any population-genetic model of heteroplasmy would assume that the germ cells or somatic stem cells giving rise to heteroplasmic variation would compete with one another for reproduction or at least be chosen randomly for transmission or reproduction. If instead, for example, there exists a cellular mechanism of quality control, such that non-heteroplasmic eggs are given priority in ovulation and tend to be ovulated before heteroplasmic eggs, the number of transmitted heteroplasmies would increase with mother’s age, but the dynamics would not be completely described by any population-genetic model that assumes random mating (with or without natural selection) and competition amongst egg cells for offspring. Other such mechanisms of heteroplasmy propagation could be imagined. Even if standard population-genetic models cannot adequately describe heteroplasmy frequency change, modeling heteroplasmy frequency changes on an ontogenetic phylogeny would still be a valid approach.

We assume that the shape of the ontogenetic phylogeny relating the sampled tissues is known. For the dataset from Rebolledo-Jaramillo *et al.* (2014), this is an appropriate assumption, since the two somatic tissues in the mother must be most closely related to one another, just as the two somatic tissues of the offspring must be most closely related to one another. For other datasets, differing in the number or identity of the sampled tissues, there may be less of an a priori expectation for the shape of the ontogenetic phylogeny. While there is a general understanding of the major divisions of tissues during development, the embryonic origins and lineage of somatic germ cell populations are not straightforward and still being established (e.g., Romagnani *et al.*, 2015; Fuentealba *et al.*, 2015; Boisset and Robin, 2012). The current model could easily be extended to ontogenetic phylogenies for families with two or more offspring. For families with more than two offspring, the genealogy of the oogonia eventually giving rise to the offspring would be unknown. This part of the phylogeny could be inferred jointly with other parameters, or, depending on the inferred rate of genetic drift in the female germ lineage (here 1.6 × 10^−3^ drift units per year), it could be assumed that no genetic drift occurs between the birth of the youngest and oldest children.

The topology of the ontogenetic phylogeny could also be made more complicated by admixture, which is not included in our inference framework. Admixture could result from biological processes, such as contributions to a mitotic tissue from distinct, isolated adult stem cell niches, or from physical sampling of an organ containing multiple tissues derived from distinct developmental lineages. Conceptually, our ontogenetic phylogeny approach could be extended to work with admixture graphs (Patterson *et al.*, 2012; Pickrell and Pritchard, 2012) by adapting the pruning algorithm for calculating likelihoods to the dependence structure introduced by admixture. However, given the small size of current heteroplasmy frequency datasets compared to large whole-genome SNP datasets, detecting admixture with f-statistics (Patterson *et al.*, 2012; Peter, 2016) or a more typical population phylogeny inference procedure (e.g., Treemix, Pickrell and Pritchard, 2012) would likely be more suitable.

The inference framework we present here should be applicable in future studies of heteroplasmy dynamics in humans and other organisms. Our software mope is flexible with respect to the pedigree of the sampled individuals and thus is suitable for studies of heteroplasmy both across several generations and within unrelated individuals. Flexibility is also given with respect to the number of tissues sampled—even studies of just a single tissue may benefit from modeling multiple ontogenetic processes (e.g., Li *et al.*, 2016). Our fully Bayesian inference method provides a natural way of quantifying uncertainty, which is important in studies of heteroplasmy as the number of polymorphic loci is often small compared to other genomic studies. Finally, mope allows the user to choose the ontogenetic processes to place in the ontogenetic phylogeny; in the current version allele frequency changes for each such ontogenetic process occur according to the neutral Wright-Fisher model, but processes governed by other dynamics (e.g., selection, mutation) could be implemented by modifying the freely available source code.

The ontogenetic phylogeny framework may also be useful in areas other than the study of mitochondrial heteroplasmy. In particular, in the study of the dynamics of cancer evolution, heterogeneous progression in samples of many tumors may necessitate modeling per-day rates of genetic drift and mutation (or natural selection) rather than fixed amounts common to all tumors. Our inference procedure could also be used in the typical population phylogenetic setting to infer the divergence history of a group of populations, but this application is limited by the relatively small number of loci (< *O*(1000)) that our method can accept due to the computational costs of likelihood evaluations with the pruning algorithm. A maximum-likelihood implementation of our model, requiring fewer likelihood evaluations, may be applicable to genome-scale SNP data, possibly comparing to Kim Tree (Gautier and Vitalis, 2013) and SpikeyTree (Tataru *et al.*, 2015).

## 5. Acknowledgments

We thank members of the Nielsen and Makova Labs for helpful comments. Two anonymous reviewers provided comments that greatly improved the manuscript. Computational resources were provided by UC Berkeley High Performance Computing. This work was funded by NIH R01GM116044.

## Appendix A. Likelihood calculation

Briefly, the pruning algorithm calculates, for each node *n* in the phylogeny and each frequency *f*_*j*_ at node *n*, the probability P(*D*_*(n)*_ | *x*_*(n)*_ = *f*_*j*_), where *D*_*(n)*_ is the data at all the leaves collectively having n as their most recent common ancestor, and *x*_*(n)*_ is the heteroplasmy allele frequency at node *n*. The algorithm proceeds up the tree, from the leaves to the root, using the fact that

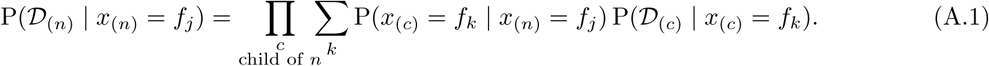

The probability P(*x*_*(c)*_ = *f*_*k*_ | *x*_*(n)*_ = *f*_*j*_) is the probability of transitioning from allele frequency *f*_*j*_ in node (*n*) to *f*_*k*_ in node (*c*), a child of (*n*). This probability is calculated using the discrete-generation Wright-Fisher model, as explained in Appendix B.

Here and below the current genetic drift parameters *b* and mutation rates *θ* are implied. We model the probability of the data at leaf (i.e., sampled tissue) node l as the binomial likelihood

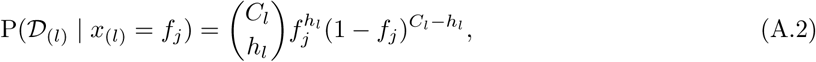

where *C*_*l*_ and *h*_*l*_ are respectively the total coverage and number of alternative alleles in that tissue. Given each P(*D*_(*r*)_ | *x*_*(r)*_ = *f*_*j*_) for root node *r*, the overall likelihood is

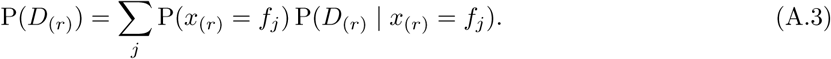

The probabilities P(*x*_*r*_ = *f*_*j*_) are given by the heteroplasmic allele frequency distribution at the root, a discretized symmetric beta distribution with additional weight at frequencies 0 and 1, the parameters of which are inferred jointly with the genetic drift and mutation parameters.

The probability of heteroplasmic polymorphism (cf. denominator of Eq. (3)) can be calculated as

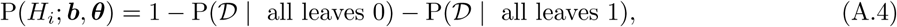

with the second two terms giving the probability of the read count data in all the sampled tissues given that allele frequencies are all 0 or 1, respectively.

## Appendix B. Calculating allele frequency transition distributions

The pruning algorithm requires distributions of allele frequency transitions along a branch. Our approach to calculating allele frequency transition probabilities is simple and intuitive: we precalculate transition distributions under the discrete-generation Wright-Fisher model using numerical matrix multiplication on a grid of generations and mutation rates. To obtain a transition distribution that was not precomputed, we linearly interpolate between precomputed distributions. Using a haploid population size of *N* = 2000 in our Wright-Fisher model calculations, we obtain a satisfactory approximation to numerically exact Wright-Fisher transition probabilities by precomputing distributions at just 207 different generations, ranging from 1 to 20, 000, and 44 mutation rates, with *θ* = 2*N*_*eμ*_ ranging from 0 to 7.5 × 10^−2^. For ontogenetic processes modeled by a single-generation bottleneck with subsequent expansion, we precompute allele-frequency transition distributions for 48 bottleneck sizes ranging from 2 to 500, linearly interpolating between bottleneck sizes for distributions that are not precomputed.

Rather than use each (2001 × 2001) transition matrix in its entirety, we combine discrete allele frequencies into 121 bins, with bins unevenly distributed between 0 and 1 such that low and high frequencies are more represented than intermediate frequencies. We bin allele frequencies according to the following scheme: Let *P* = {*P*_*ij*_} be a (2001 × 2001) allele frequency transition matrix for a Wright-Fisher model with *N* = 2000, with *P*_*ij*_ being the probability of transitioning from frequency *i* to *j*. Let *Q* = {*Q*_*k, l*_} be a (121 × 121) binned transition matrix. If (*a*_1_,…, *a*_*m*_) are frequencies in bin *k*, and (*b*_1_,…, *b*_*n*_) are frequencies associated in bin l, then

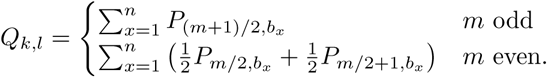

## Appendix C. Calculation of the effective bottleneck size

We define the effective bottleneck between mother and offspring as the combined genetic drift occurring during the early oogenic bottleneck, the turnover of mitochondria in the maternal germline prior to ovulation, and the first few cell divisions after fertilization but before gastrulation. We combined the effects of genetic drift during these processes by 1) translating all drift parameters into units of generations per effective population size (*g*/*N*_*e*_, “drift units”), 2) summing the drift, in these units, and 3) translating this summed drift back into units of an instantaneous bottleneck. Since we assumed that bottlenecks occurred for just a single generation followed by doubling back up to a large population size (here, *N* = 2000), we determined that the relationship between drift *d*_*g*_ measured in drift units and *N*_*b*_, an instantaneous bottleneck size, is close to

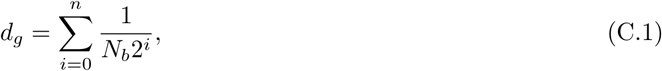

where *n* = ⌊log_2_(*N*/*N*_*b*_)⌋ is the number of generations it takes for the population size to double back up to the original population size.

For *N*_*b*_ << *N*, this sum is well approximated by the integral

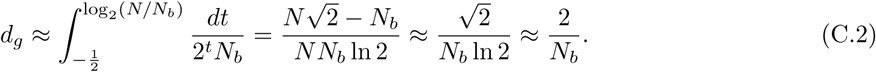

The lower limit of integration follows from an interpretation of (C.1) as a midpoint Riemann sum, improving accuracy. Thus we also have

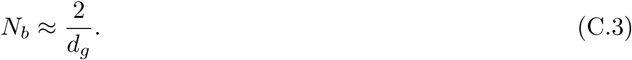

For a mother of age *a*, the effective bottleneck size is thus

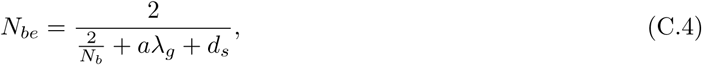

where *N*_*b*_ is the early oogenesis bottleneck size, *λ*_*g*_ is the rate at which genetic drift accumulates in the maternal germline, and *d*_*s*_ is the amount of genetic drift occurring after fertilization but before gastrulation.

We confirmed (C.2) and (C.3) by finding, for different bottleneck sizes *N*_*b*_, the amount of drift *d*_*g*_ that minimized the total variation distance between the allele frequency transition distributions specified by *d*_*g*_ and *N*_*b*_:

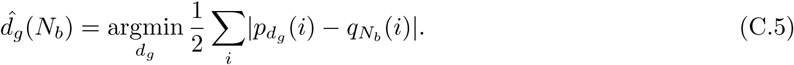

Here *p*_*dg*_ is the probability transition distribution for drift parameterized by *d_g_* drift units, and *q*_*Nb*_ is the probability transition distribution for drift parameterized by bottleneck size *Nb*. Minimizing (C.5) for different values of *Nb* shows that our approximation (C.2) closely follows the numerically translation minimizing the total variation distance (Fig. S5).

---

**Figure S1:**
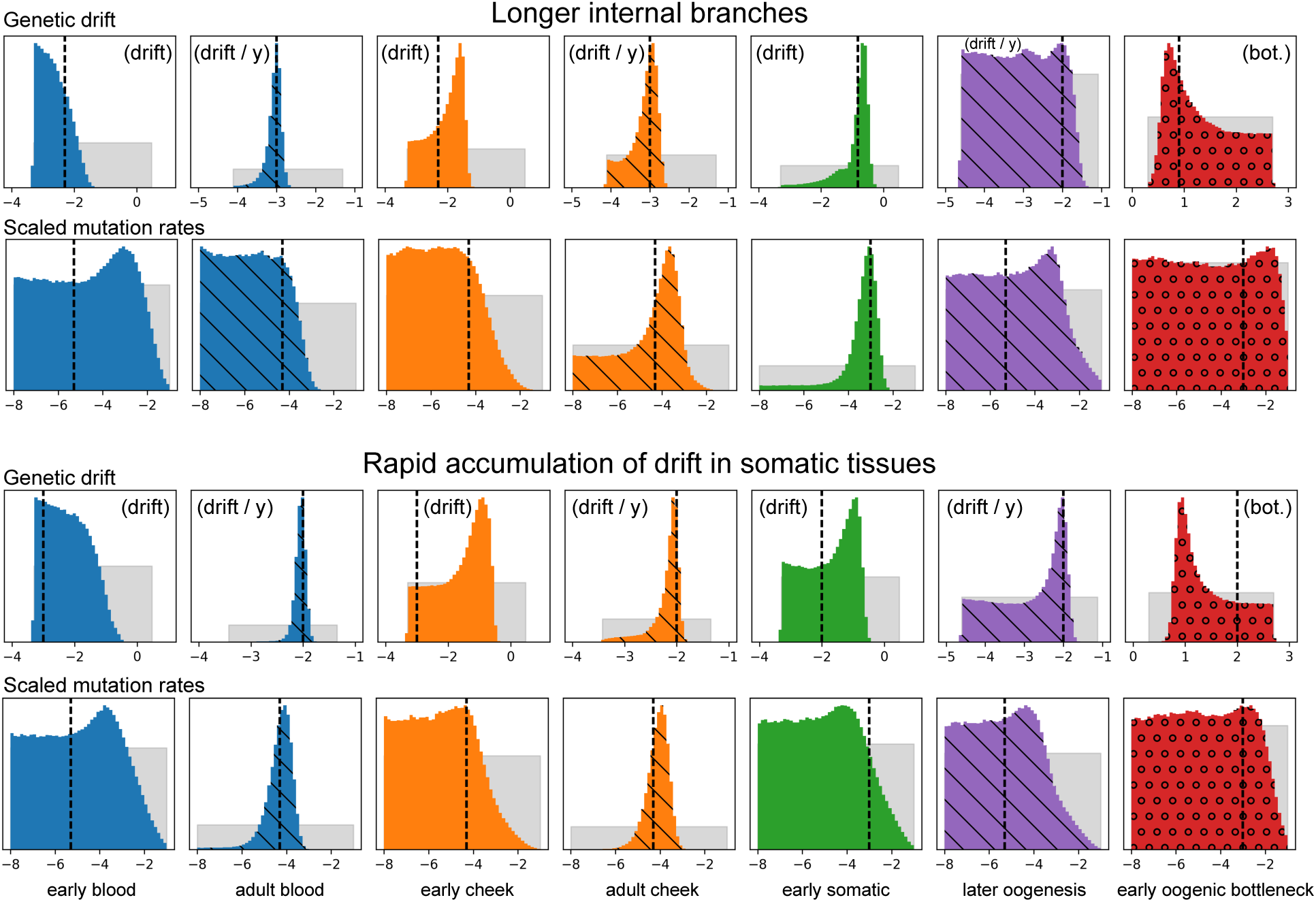
Inference results from additional simulations under the model assumed by our inference procedure. The first two rows show posterior distributions of parameters estimated under simulations in which the internal branches are relatively long compared to the simulations presented in the main text. These parameters were inferred from the frequencies of 103 hetero-plasmic loci amongst 500 independently sites in 40 simulated families. The second pair of rows shows posterior distributions for simulations in which the rates of accumulation of genetic drift in the somatic tissues is increased compared to the simulations in the main text. In these simulations, there were 109 heteroplasmic loci amongst 400 independently segregating loci simulated in 80 families. Posterior distributions are shown with colored histograms, prior distributions are shown with gray histograms, and true parameter values are shown with dashed vertical lines. Colors match the corresponding developmental processes in Figure 1. Distributions hashed with diagonal lines correspond to processes with drift parameterized by rates of accumulation of genetic drift with age, and circles in the red posterior distributions indicate that this process is modeled by an explicit bottleneck.

**Figure S2:**
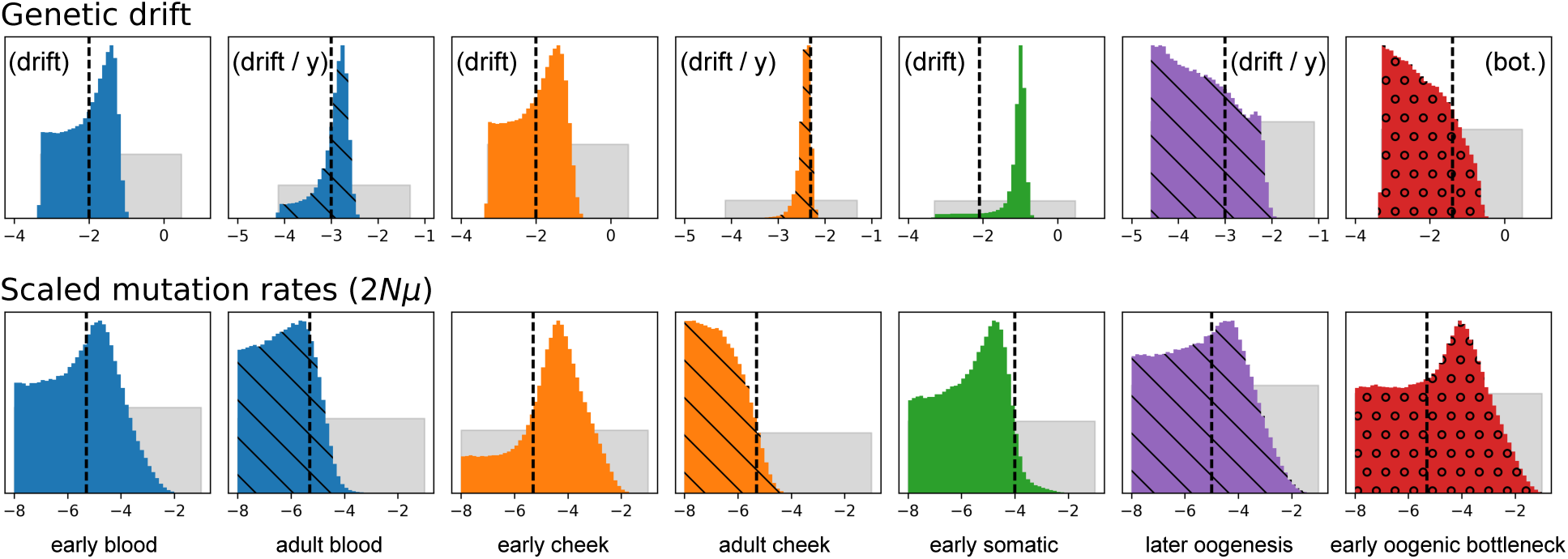
Inference results from simulations with no recombination between heteroplasmic loci segregating within a single family. The first row shows posterior distributions (color histograms), prior distributions (gray distributions) and simulated parameter values (dashed vertical lines) for genetic drift parameters. The second row shows the same for scaled mutation rate parameters. In order to simplify simulations, the period of genetic drift in early oogenesis was modeled as a period of genetic drift in a fixed population size rather than as a single-generation bottleneck.

**Figure S3:**
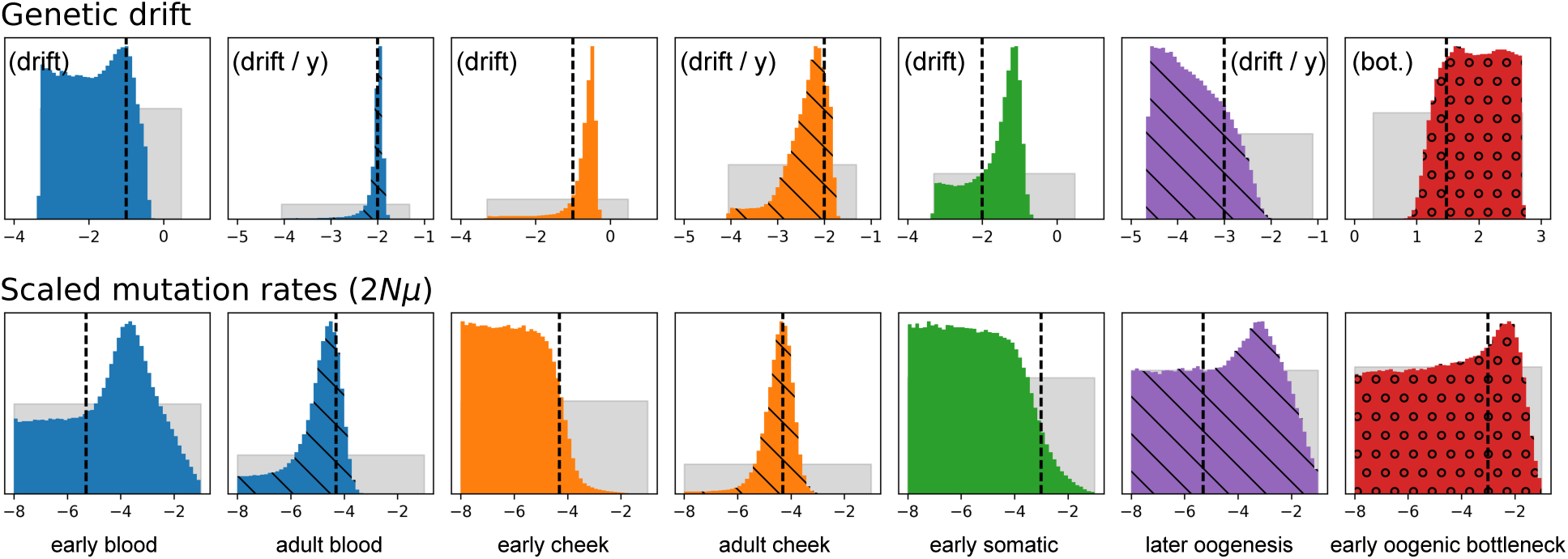
Inference results from simulations with noisy heteroplasmy detection. False negatives were simulated with probability 0.4 for each truly heteroplasmic locus with frequency between 0.1% and 2%. False negatives were produced at a rate of 3×10^−5^ per bp, so that 14 false negatives and 5 false positives were produced in a dataset of 101 heteroplasmic loci. As in other figures presented here, the first row shows posterior distributions (color histograms), prior distributions (gray distributions) and simulated parameter values (dashed vertical lines) for genetic drift parameters. The second row shows the same for scaled mutation rate parameters.

**Figure S4:**
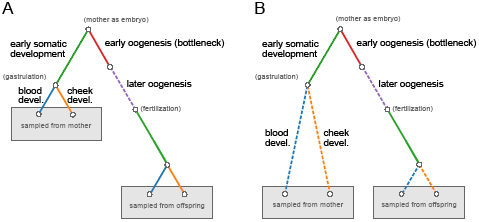
Two additional ontogenetic phylogenies for which we calculated the total Bayesian evidence. The two models differ in how they model genetic drift and mutation in the somatic tissues. The “fixed” model (Panel A) assumes that all genetic drift and mutation in the somatic tissues occurs early in development, and the the “linear” model (Panel B) assumes that genetic drift and mutation in somatic tissues accumulate linearly with the age of the individual. Compare these to the model in Figure 1 (termed “both”), which assumes that genetic drift and mutation in somatic tissues occurs both in a fixed amount during early development and in adulthood, accumulating linearly with age.

**Figure S5:**
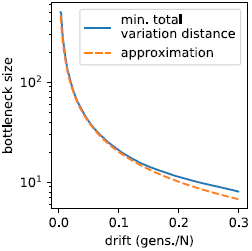
Translation of genetic drift into effective bottleneck sizes. The blue line shows, for different drift durations, the effective bottleneck size minimizing the total variation distance to the allele frequency transition distribution parameterized by generations per effective population size. The dashed orange line shows Equation (C.3), our approximate translation between the two parameterizations.

## References

Boisset, J.-C., and C. Robin, 2012 On the origin of hematopoietic stem cells: Progress and controversy. Stem Cell Research 8: 1–13.

Burgstaller, J., I. Johnston, N. Jones, J. AlbrechtovaÁ, T. Kolbe, et al., 2014 mtDNA segregation in heteroplasmic tissues is commin in vivo and modulated by haplotype differences and developmental stage. Cell Reports 7: 2031–2041.

Cao, L., H. Shitara, T. Horii, Y. Nagao, H. Imai, et al., 2007 The mitochondrial bottleneck occurs without reduction of mtDNA content in female mouse germ cells. Nature Genetics 39: 386–390.

Carling, P. J., L. M. Cree, and P. F. Chinnery, 2011 The implications of mitochondrial DNA copy number regulation during embryogenesis. Mitochondrion 11: 686–692.

Carlson, B. M., 2014 Human Embryology and Developmental Biology. Elsevier, Philadelphia, 5 edition.

Cree, L. M., D. C. Samuels, S. C. De Sousa Lopes, H. K. Rajasimha, P. Wonnapinij, et al., 2008 A reduction of mito-chondrial DNA molecules during embryogenesis explains the rapid segregation of genotypes. Nature Genetics 40: 249–254.

Ewens, W. J., 2004 Mathematical Population Genetics 1. Number 27 in Interdisciplinary Applied Mathematics. Springer, New York, 2 edition.

Fan, W., K. G. Waymire, N. Narula, P. Li, C. Rocher, et al., 2008 A mouse model of mitochondrial disease reveals germline selection against severe mtDNA mutations. Science 319: 958–962.

Felsenstein, J., 1981 Evolutionary trees from DNA sequences: a maximum likelihood approach. Journal of Molecular Evolution 17: 368–376.

Foreman-Mackey, D., D. W. Hogg, D. Lang, and J. Goodman, 2013 emcee: The MCMC hammer. Publications of the Astronomical Society of the Pacific 125: 306–312. ArXiv: 1202.3665.

Fuentealba, L., S. Rompani, J. Parraguez, K. Obernier, R. Romero, et al., 2015 Embryonic origin of postnatal neural stem cells. Cell 161: 1644–1655.

Gautier, M., and R. Vitalis, 2013 Inferring population histories using genome-wide allele frequency data. Molecular Biology and Evolution 30: 654–668.

Goggans, P. M., and Y. Chi, 2004 Using thermodynamic integration to calculate the posterior probability in Bayesian model selection problems. AIP Conference Proceedings 707: 59–66.

Goodman, J., and J. Weare, 2010 Ensemble samplers with affine invariance. Communications in Applied Mathematics and Computational Science 5: 65–80.

Guo, Y., C.-I. Li, Q. Sheng, J. F. Winther, Q. Cai, et al., 2013 Very low-level heteroplasmy mtDNA variations are inherited in humans. Journal of genetics and genomics 40: 607–615.

Hendy, M. D., M. D. Woodhams, and A. Dodd, 2009 Modelling mitochondrial site polymorphisms to infer the number of segregating units and mutation rate. Biology Letters: rsbl.2009.0104.

Jenuth, J. P., A. C. Peterson, K. Fu, and E. A. Shoubridge, 1996 Random genetic drift in the female germline explains the rapid segregation of mammalian mitochondrial DNA. Nature Genetics 14: 146–151.

Johnston, I. G., J. P. Burgstaller, V. Havlicek, T. Kolbe, T. Rlicke, et al., 2015 Stochastic modelling, Bayesian inference, and new in vivo measurements elucidate the debated mtDNA bottleneck mechanism. eLife 4: e07464.

Johnston, I. G., and N. S. Jones, 2016 Evolution of cell-to-cell variability in stochastic, controlled, heteroplasmic mtDNA populations. The American Journal of Human Genetics 99: 1150–1162.

Kelleher, J., A. M. Etheridge, and G. McVean, 2016 Efficient coalescent simulation and genealogical analysis for large sample sizes. PLOS Comput Biol 12: e1004842.

Khrapko, K., 2008 Two ways to make an mtDNA bottleneck. Nature Genetics 40: 134.

Li, M., R. Rothwell, M. Vermaat, M. Wachsmuth, R. Schrder, et al., 2016 Transmission of human mtDNA heteroplasmy in the Genome of the Netherlands families: support for a variable-size bottleneck. Genome Research 26: 417–426.

Li, M., A. Schönberg, M. Schaefer, R. Schroeder, I. Nasidze, et al., 2010 Detecting heteroplasmy from high-throughput sequencing of complete mitochondrial DNA genomes. The American Journal of Human Genetics 87: 237–249.

Li, M., R. Schr oder, S. Ni, B. Madea, and M. Stoneking, 2015 Extensive tissue-related and allele-related mtDNA hetero-plasmy suggests positive selection for somatic mutations. Proceedings of the National Academy of Sciences 112: 2491–2496.

Li, M., and M. Stoneking, 2012 A new approach for detecting low-level mutations in next-generation sequence data. Genome Biology 13: R34.

Marchington, D. R., G. M. Hartshorne, D. Barlow, and J. Poulton, 1997 Homopolymeric tract heteroplasmy in mtDNA from tissues and single oocytes: support for a genetic bottleneck. American Journal of Human Genetics 60: 408–416.

Millar, C. D., A. Dodd, J. Anderson, G. C. Gibb, P. A. Ritchie, et al., 2008 Mutation and evolutionary rates in Adélie penguins from the Antarctic. PLoS Genetics 4: e1000209.

Patterson, N., P. Moorjani, Y. Luo, S. Mallick, N. Rohland, et al., 2012 Ancient admixture in human history. Genetics 192: 1065–1093.

Peter, B. M., 2016 Admixture, population structure, and f-statistics. Genetics 202: 1485–1501.

Pickrell, J. K., and J. K. Pritchard, 2012 Inference of population splits and mixtures from genome-wide allele frequency data. PLoS Genetics 8: e1002967.

Rebolledo-Jaramillo, B., M. S.-W. Su, N. Stoler, J. A. McElhoe, B. Dickins, et al., 2014 Maternal age effect and severe germ-line bottleneck in the inheritance of human mitochondrial DNA. Proceedings of the National Academy of Sciences 111: 15474–15479.

Romagnani, P., Y. Rinkevich, and B. Dekel, 2015 The use of lineage tracing to study kidney injury and regeneration. Nature Reviews Nephrology 11: 420–431.

Samuels, D. C., C. Li, B. Li, Z. Song, E. Torstenson, et al., 2013 Recurrent tissue-specific mtDNA mutations are common in humans. PLOS Genetics 9: e1003929.

Sharpley, M., C. Marciniak, K. Eckel-Mahan, M. McManus, M. Crimi, et al., 2012 Heteroplasmy of mouse mtDNA is genetically unstable and results in altered behavior and cognition. Cell 151: 333–343.

Sondheimer, N., C. E. Glatz, J. E. Tirone, M. A. Deardorff, A. M. Krieger, et al., 2011 Neutral mitochondrial hetero-plasmy and the influence of aging. Human Molecular Genetics 20: 1653–1659.

Stewart, J. B., and P. F. Chinnery, 2015 The dynamics of mitochondrial DNA heteroplasmy: implications for human health and disease. Nature Reviews Genetics 16: 530–542.

Stewart, J. B., C. Freyer, J. L. Elson, A. Wredenberg, Z. Cansu, et al., 2008 Strong purifying selection in transmission of mammalian mitochondrial DNA. PLOS Biology 6: e10.

Tataru, P., T. Bataillon, and A. Hobolth, 2015 Inference under a Wright-Fisher model using an accurate beta approximation. Genetics 201: 1133–1141.

Wachsmuth, M., A. Hubner, M. Li, B. Madea, and M. Stoneking, 2016 Age-related and heteroplasmy-related variation in human mtDNA copy number. PLoS Genetics 12: e1005939.

Wai, T., D. Teoli, and E. A. Shoubridge, 2008 The mitochondrial DNA genetic bottleneck results from replication of a subpopulation of genomes. Nature Genetics 40: 1484–1488.

Wakeley, J., 2009 Coalescent Theory: An Introduction. Roberts and Co., Greenwood Village, CO, 1 edition.

Wallace, D. C., and D. Chalkia, 2013 Mitochondrial DNA genetics and the heteroplasmy conundrum in evolution and disease. Cold Spring Harbor Perspectives in Biology 5: a021220.

Wonnapinij, P., P. F. Chinnery, and D. C. Samuels, 2008 The distribution of mitochondrial DNA heteroplasmy due to random genetic drift. The American Journal of Human Genetics 83: 582–593.

Ye, K., J. Lu, F. Ma, A. Keinan, and Z. Gu, 2014 Extensive pathogenicity of mitochondrial heteroplasmy in healthy human individuals. Proceedings of the National Academy of Sciences 111: 10654–10659.

Zhang, D., D. Keilty, Z. Zhang, and R. Chian, 2017 Mitochondria in oocyte aging: current understanding. Facts, Views & Vision in ObGyn 9: 29–38.

